# Speed-Selectivity in Retinal Ganglion Cells is Modulated by the Complexity of the Visual Stimulus

**DOI:** 10.1101/350330

**Authors:** César R Ravello, Laurent U Perrinet, María-José Escobar, Adrián G Palacios

## Abstract

Motion detection represents one of the critical tasks of the visual system and has motivated a large body of research. However, is remain unclear precisely why the response of retinal ganglion cells (RGCs) to simple artificial stimuli does not predict their response to complex naturalistic stimuli. To explore this topic, we use Motion Clouds (MC), which are synthetic textures that preserve properties of natural images and are merely parameterized, in particular by modulating the spatiotemporal spectrum complexity of the stimulus by adjusting the frequency bandwidths. By stimulating the retina of the diurnal rodent, *Octodon degus* with MC we show that the RGCs respond to increasingly complex stimuli by narrowing their adjustment curves in response to movement. At the level of the population, complex stimuli produce a sparser code while preserving movement information; therefore, the stimuli are encoded more efficiently. Interestingly, these properties were observed throughout different populations of RGCs. Thus, our results reveal that the response at the level of RGCs is modulated by the naturalness of the stimulus - in particular for motion - which suggests that the tuning to the statistics of natural images already emerges at the level of the retina.

## Introduction

Motion detection is essential for animal survival, and studies show that many retinal cells^1–5^ and a considerable portion of the cortex^6,7^ are involved in its processing. However, most of these studies have focused on the detection of artificial stimuli, such as random dots, lines, or moving gratings, while ignoring the natural signals from the environment in which animals have evolved and to which their visual system would be attuned to^8–13^. Moreover, computational models based on the response to simple, artificial stimuli fail to predict the response to naturalistic images^14–16^. Nevertheless, working with natural images directly is not always possible due to their complexity regarding critical parameters of signals processing, including visual content and its variability. The latter makes it hard to control the motion information that is effectively present to the visual system^17,18^.

In the visual cortex, the motion is detected at different levels and by populations of cells attuned to different parameters of the stimulus (see^6,7^ for reviews). It has been reported that visual neurons stimulated with complex images, which should be closer to natural images, change their tuning curves compared to the response obtained with simple stimuli, becoming narrower and thus more selective to speed^19^. In turn, stimulation with naturalistic images generates sparser code^20–22^ and eye tracking responses become more precise^23^.

In the retina, the most basic form of motion detection is the response to moving textures in which Y-like *Retinal Ganglion Cells* (RGCs) increase their firing rate when sub-regions of their *Receptive Field* (RF) alternate between light and dark luminosity levels through time, regardless of the direction of motion^24–26^. It has been demonstrated that these mechanism arises from the RF properties of ON and OFF *Bipolar Cells* (BCs) that connect to the RGCs^27^ and transmit their activity in a non-linear fashion. This non-linear integration of the activity of BCs allows that the alternating activation of cells of both polarities do not cancel each other upon reaching RGCs^28^. These BCs can generate fast transient responses when there is a luminance change of the corresponding polarity (dark to light or light to dark respectively), so each time that BCs detects a temporal change in luminance, it will generates an EPSPs in the corresponding RGCs. The integration time of the RGCs is longer than the time course of the response of the BCs, so fast successive activation will be added. In this way, the RGCs will respond with higher firing rates to higher speeds because of the higher rate of alternating activation of the ON and OFF sub-fields.

More specialized computations are carried out by, e.g., *Direction Selective* RGCs which respond preferentially to a single direction in a relatively narrow range of speed^1,5,29,30^, with different subtypes tuned to different directions oriented perpendicularly, and also with different polarities; *Object Motion* RGCs detectors, which fire when a small patch of the image moves in a different pattern than the rest of the background^4,31,32^ and *Approaching Motion* RGCs detectors which fire when a dark patch increases its size through time^33^. Another sophisticated processing of motion is the response to *Motion Onset* RGCs detectors ^34^, in which the initiation of motion within the RF causes a higher response than does an object moving smoothly through it. While it has been shown that the retina is capable of advanced computations beyond standard processing^35^, many capabilities remain unexplored^36^, for example the fine tuning of retinal responses to motion in natural images.

In this study, we focus on the properties of RGCs responses to motion in several scenarios. To overcome the limitations of working with simple, low-dimensional artificial stimuli, while at the same time avoiding the problems of working with natural images, we used synthetic random textures called Motion Clouds (MC)^37^, which preserve some of the properties of natural images. MC they mimic the spatiotemporal changes in luminance of natural scenes^38^. They correspond to a mix of moving gratings with random positions and spatiotemporal properties distributed along a parameterizable area (in Fourier space), *i.e*. the components will have different sizes (given by their spatial frequency) and will move at different speeds (provided by their slope in the spatiotemporal plane) in contrast to traditional drifting grating which has a single spatial and temporal frequency; (see Figure 1a for comparison of these two stimuli). While natural images have their power distributed along a broad range of frequencies, MC can be parametrized to have a restricted range of frequencies to limit their information content or complexity and parameterized by the bandwidth parameter (see Figure 1b). They have been used successfully to study speed discrimination in humans^23^, where it was shown that additional information contained in the stimuli improves eye pursuit gain and precision.

**Figure 1.**
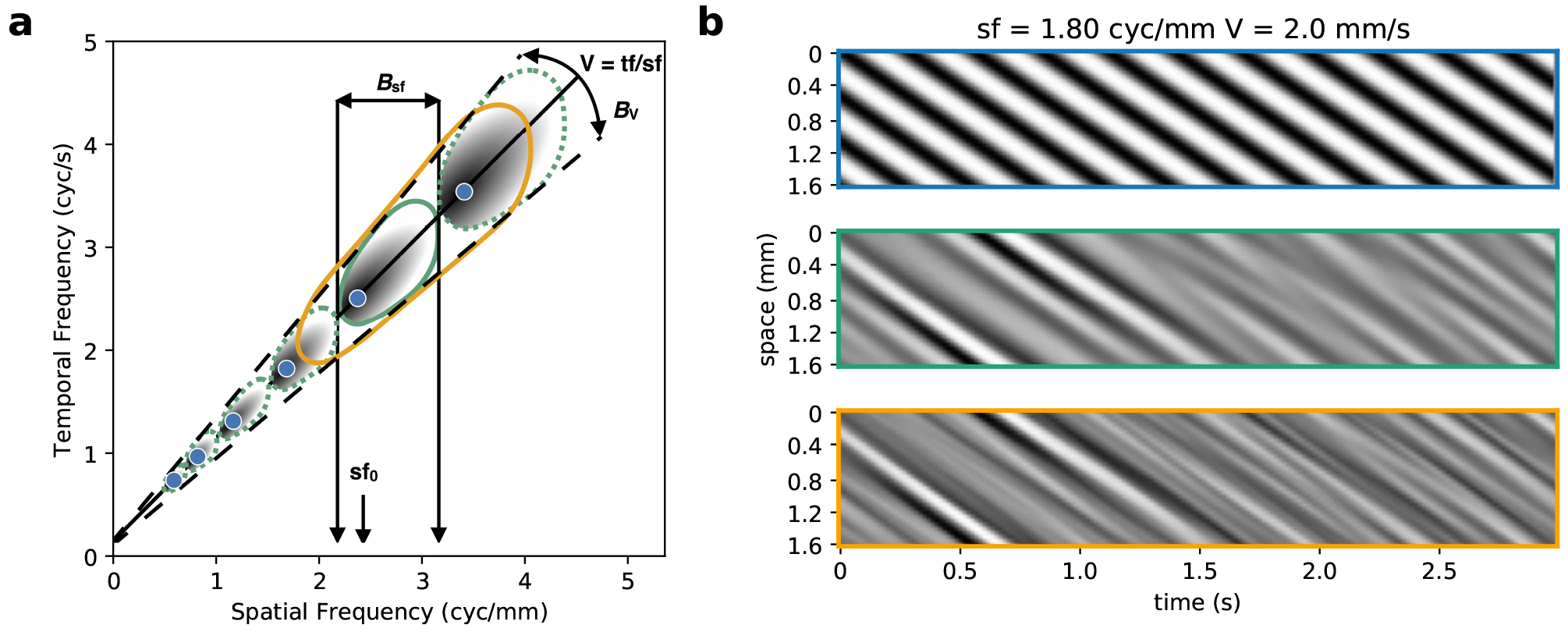
Motion Cloud (MC) stimuli are characterized by the parameters of their spatio-temporal spectral envelope. **a** Bi-dimensional representation of the spatio-temporal frequency space. Points along the continuous line correspond to simple drifting grating stimuli moving at a given speed but with different spatial frequencies (blue dots). MC stimuli have their spectral energy distributed in an “ellipse” warped along a given speed plane. Interestingly, these can be parameterized with the same mean spatial (sf_0_) frequency and speed (*V*) as gratings, but with different levels of spatial frequency bandwidths (defined by the parameter *B*_sf_, with *B*_sf_ = 0 for the grating) which will define the level of complexity of the sequence. The green ellipse represents a narrow bandwidth stimulus and the orange ellipse a broader bandwidth stimulus. **b** S Examples of stimuli at different complexity levels. For simplicity, we plot the light intensity along a single row of the image (vertical axis) for the whole duration of the stimulation (horizontal axis). We show the drifting grating (top panel), which is constituted by a single spatial and temporal frequency (and thus a single speed, seen as the slope in this view), and the MCs (middle and lower panels), which have the same speed and central spatial frequency but with a narrow or wide bandwidth in spatial frequency space.

**Figure 2.**
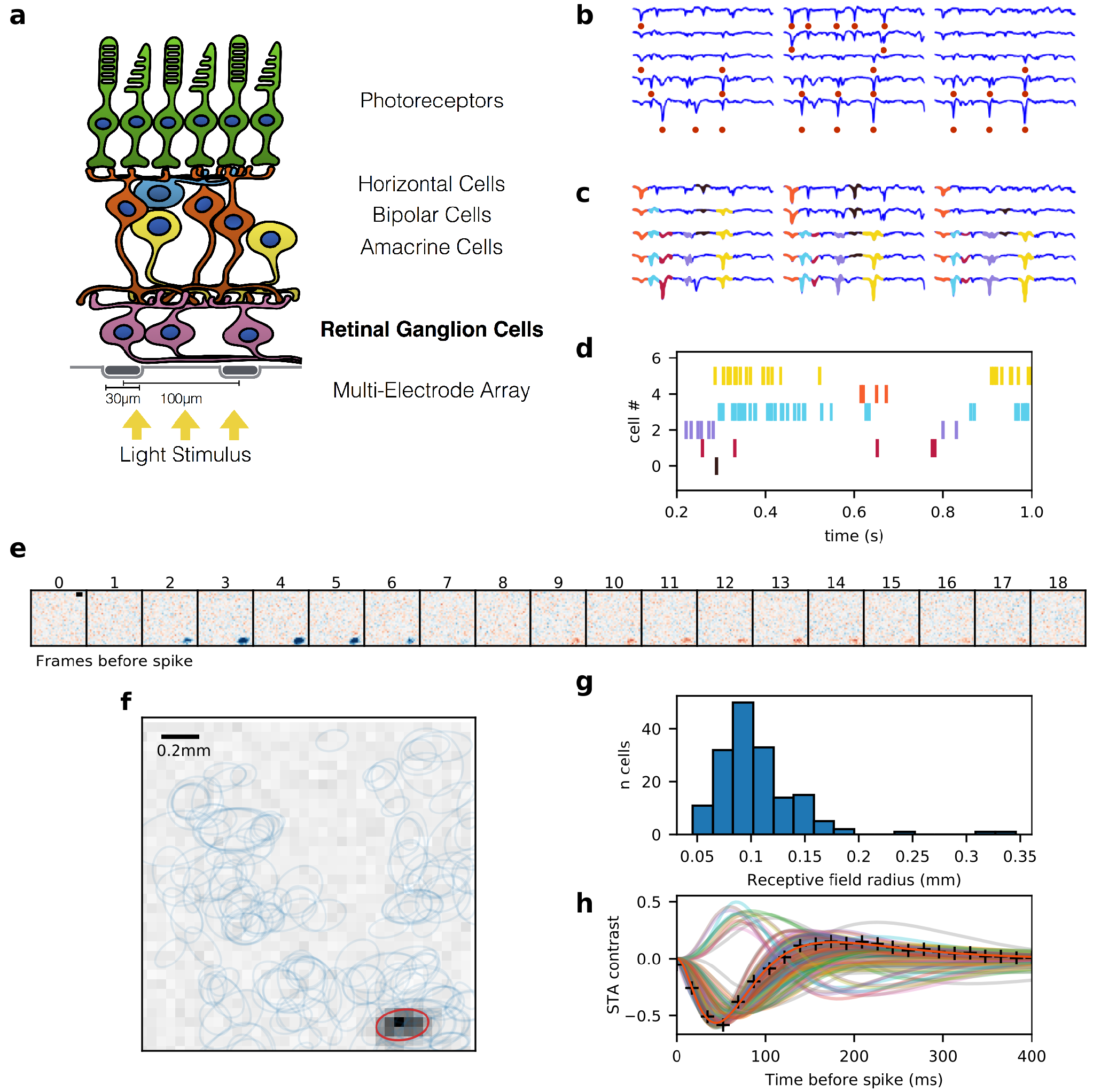
Experimental multielectrode recording setup. **a** Schematic view of the retina over a Multielectrode Array, while light stimulation is projected from below. **b** Each blue line represents a voltage signal from an electrode, and the red dots indicate detected peaks. **c** The signal of each electrode is reconstructed by iteratively adding different templates. Each color represents a different template, indicating spikes coming from different cells. Each spike can be detected by many neighboring electrodes, but only one will be assigned to a single unit. **d** The raster plot indicates the times at which each cell fired an action potential. **e** Spike Triggered Average (STA) of a representative cell, computed from the response to checkerboard stimulus. Each panel shows the average image at the corresponding frame before the spiking of the cell (time zero). Blue pixels represent points with negative contrast, while red represent those with positive contrast. The black bar is 0.2 mm long. **f** Spatial component of the STAs of all the cells recorded from a retinal patch, after discarding non-valid units (see Methods for details). Each ellipse corresponds to a 2-D Gaussian fit at 1 s.d. The background shows the frame from (e) of maximum response and the red ellipse its fit, while the blue ellipses show the fit for the rest of the cells. **g** Distribution of RF sizes from a retinal patch, measured as the radius of a circle with the same area as the ellipse fit. **h** Temporal components of the STA of the cells shown in (g). The crosses mark the STA of the cell shown in (e).

The activity recorded from a population of RGCs was compared using simple moving stimuli (drifting gratings) vs. MC’s with the same speed and two levels of naturalness, regarding their spatial frequency spectrum (narrow and broad bandwidth). We found that for a significant fraction of RGCs responding to the motion, the cells’ tuning bandwidths become narrower when the spatiotemporal frequency content of the stimulus increased. The set of RGCs studied contains cells traditionally assigned to different types, indicating that this property is not exclusive of a single class, but instead is. At the population level, this change in tunning bandwidth results in sparser codes, which in turn reflects a more efficient coding of the motion information. Therefore, our findings show that fine-tuning of motion detection to natural image statistics already emerges at the level of the retina.

## Results

We characterized the response from a population of *speed responsive* RGCs, recorded from retinal patches of 2 young *Octodon degus*, a diurnal rodent, having a total of 308 RGCs (see Methods section for details).

### Speed responsive cells in the retina as characterized using gratings

We measured RGCs responses to a set of drifting gratings (artificial simple stimuli), with varying speed and spatial frequency (see Figure 3a). For each spatial frequency, the response to speed averaged over trials was fitted to a skewed Gaussian. In Figure 3b, we show how preference for certain speeds changes as a function of spatial frequency. As expected for RGCs, preferred speed decreases at higher spatial frequencies. In the example shown, the response is maximal at intermediate speeds and spatial frequencies, diminishing gradually as one moves away from that preferred combination of parameters. When looking at the fitted curves, it can be seen that the whole curve shifts towards lower speeds when increasing the spatial frequency.

**Figure 3.**
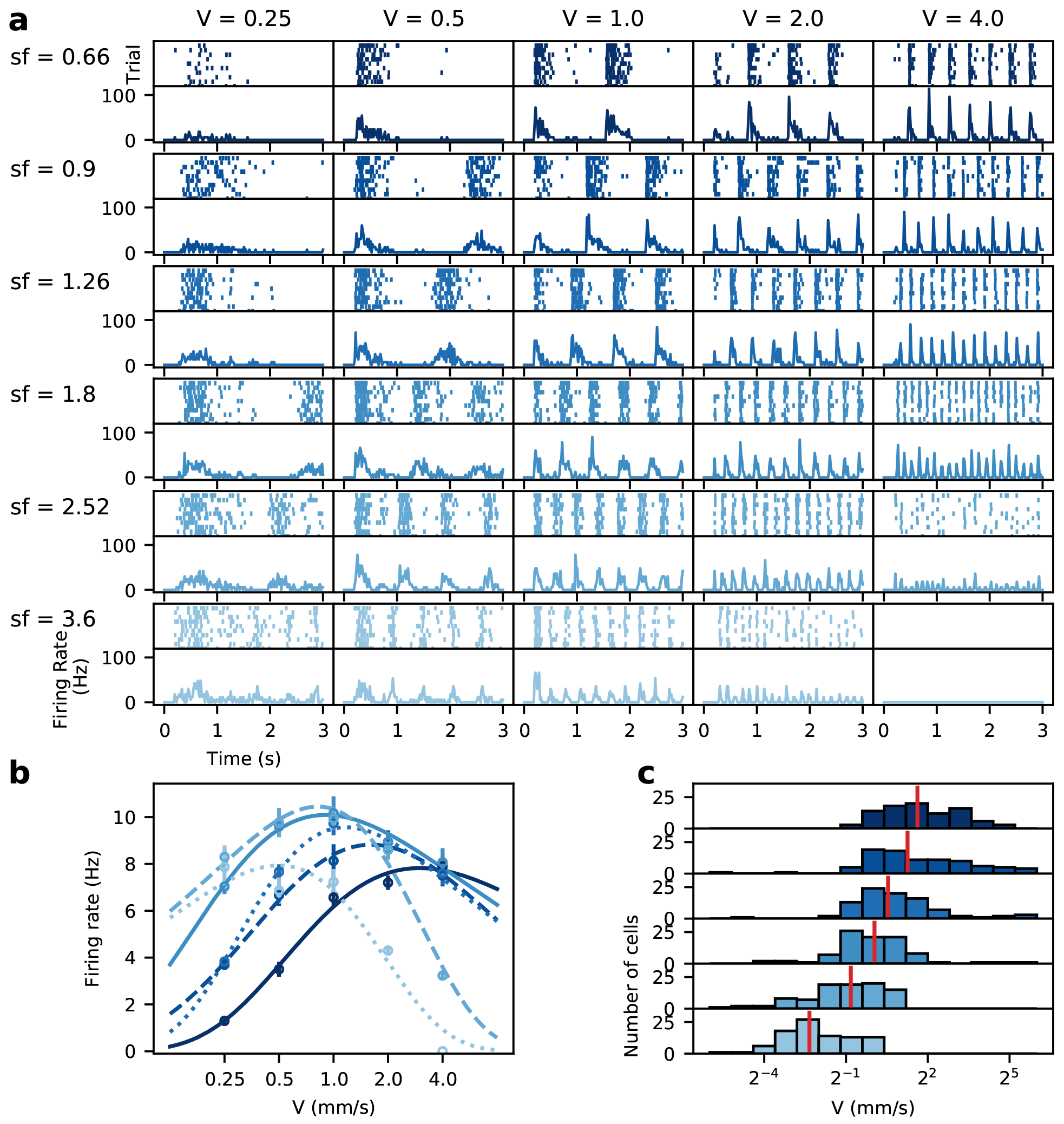
Response to variations in speed and spatial frequency of the drifting grating stimulus. **a** Raster and PSTHs of the response of a representative cell at each speed and spatial frequency tested. **b** The response to speed at each spatial frequency sf was fitted to a skewed Gaussian. Each point is the average of the PSTH, with the error bars showing SEM. **c** All cells that had a good fit (*χ*^2^ < 0.05) at every sf were classified as *speed responsive cells*. For all these cells, the preferred speed at each spatial frequency is determined from the fits to their distribution. Vertical red lines show the median of the distribution for each condition.

A subset of the complete set of valid RGCs was separated and classified as *speed responsive cells* (SR), *i.e*. cells whose response to the speed of the drifting grating has a good fit to Equation 2 at every spatial frequency tested (*χ*^2^ < 0.05 for the normalized response), see Figure 3b. We analyzed retinal patches from two animals, where 144 and 164 SR cells were found, respectively, representing ≈ 46% and ≈ 44% of the total RGCs recorded, respectively. Figure 4a shows the total number of RGCs found in a sample piece of retina, and their respective RF, of which the SR cells are highlighted in color.

**Figure 4.**
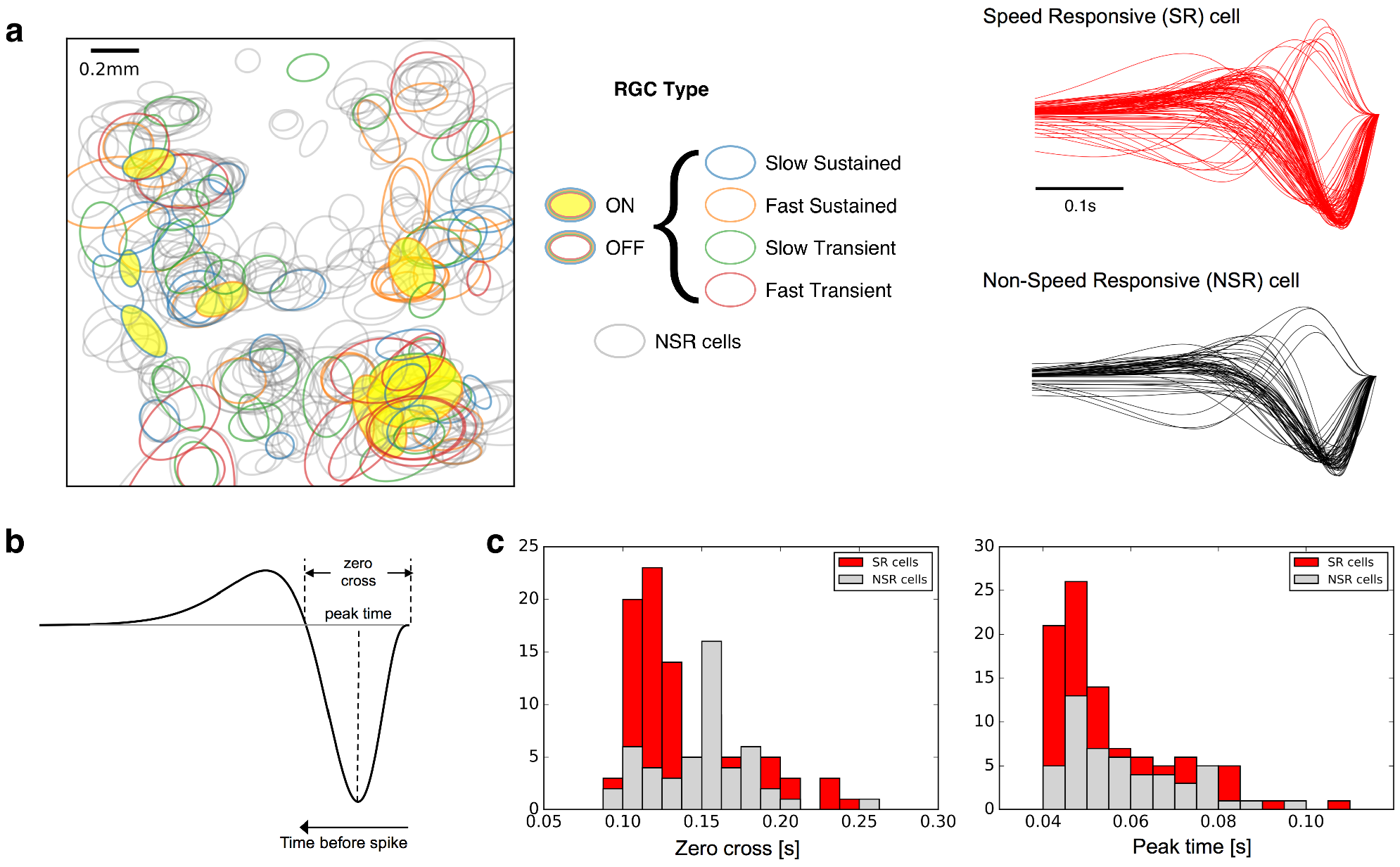
Speed responsive retinal ganglion cells. **a** *Left:* Map of the RGCs RF present in the experiment analyzed. RF of speed responsive (SR) RGCs (see text) are represented in color. *Right:* Temporal profiles of the receptive field center of SR (in red) and Non-SR (in black) cells. **b** Sample of a temporal profile showing the parameters extracted to compare SR versus Non-SR cells, which in this case are zero-cross and peak-time. **c** Histograms representing the distribution of the zero-cross (left) and peak-time (right) parameters in the SR and Non-SR population cells. We only observe significant differences for the zero-cross parameter (*p* < 0.01 versus *p* > 0.1 for the peak-time parameter, Kolmogorov-Smirnof test).

To discriminate between SR and Non-SR cells, we analyzed the RF properties of each class. The temporal profile of a cell’s response was characterized using the parameters indicated in Figure 4b: zero-cross and peak-time. In Figure 4c we show the distribution of these two parameters along the entire population of RGCs, separated by their speed responsive selectivity (SR cells are represented in red, while Non-SR in black). Interestingly, only the zero-cross parameter is significantly different between SR and Non-SR cell populations (*p* < 0.01 Kolmogorov-Smirnov test). Additionally, there was no direct correlation with other traditional classification parameters of RGCs, such as Fast-Slow and Transient-Sustained using the latency and transience index^39^ of the response to a full-field flash, since our set of SR cells contained cells belonging to each class (28 Fast-Transient, 15 Fast-Sustained, 22 Slow-Transient and 28 Slow-Sustained from 93 SR cells).

### Speed selectivity changes with the complexity of the stimulus

When presenting a stimulus with the same base properties as the drifting gratings but with energy distributed along a larger area in parameter space, the responses differ significantly. Examining in detail the response at the preferred spatial frequency (Figure 5a) reveals that the response to lower and higher speeds is much weaker for the MC stimuli compared to the response to the grating, while the responses become equivalent in terms of firing rate only around the preferred, intermediate speeds. This effect is more evident when looking at the tuning curves (Figure 5b), where the response decreases more sharply when moving away from the preferred parameters for the complex stimuli. The change in the shape of the curve is evidenced by a change in the σ value of the fit, which decreases progressively from 1.76 to 1.45 and 0.95 as the complexity of the stimulus increases, while maintaining the same preferred speed (1.47, 1.37 and 1.49 mm/ sec respectively). The firing rates are stable across repetitions, so the narrower tuning is not due to drifts or decay in cell activity. When considering all the conditions tested, the response profile to the MC stimuli cover a smaller area in parameter space along both axes, when viewing it as a two dimensional map of the response at each combination of parameters (Figure 5c). As expected, speed preference decreases with spatial frequency, as seen by the slanted response profile.

**Figure 5.**
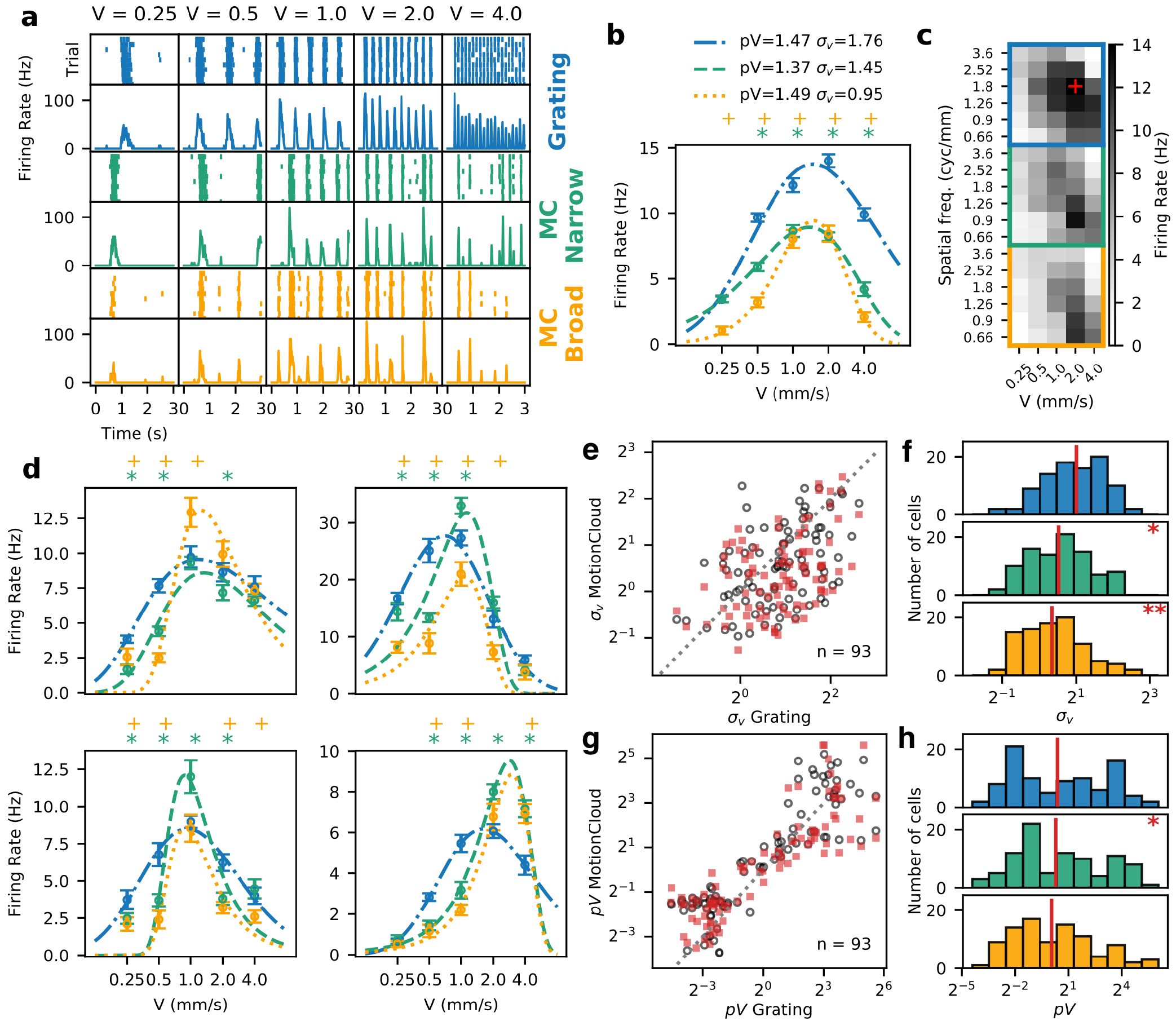
A large part of the population of observed RGCs narrow down their tuning selectivity with higler complexity stimuli. **a** Raster and PSTHs of the response of a representative cell for each type of stimulus and to the different speeds (in mms^−1^) at its preferred sf_0_. **b** The change in tuning is evident when plotting the average response and its fit. Each point is the average of the PSTH, with the error bars showing s.d. The asterisks (for the Narrow bandwidth) and crosses (Broad bandwidth) mark the speeds at which the response to the MC is significantly different from the response to the grating (*p* < 0.05 Wilcoxon signed-rank test). The narrowing of the curve can be quantified by the decrease in the *σ*_*v*_ parameter, which in the case of this cell decreased with the complexity of the stimulus. **c** Spatiotemporal tuning of the cell for the different types of stimuli. The intensity at each square denotes the average firing rate for the corresponding stimulus parameters. The red cross indicates the point of maximum response, and corresponds to the spatial frequency plotted in panels a and b. **d** Additional examples of cells with narrower tuning for the naturalistic stimulus. **e** Joint distribution of speed tuning bandwidth coefficients *σ*_*v*_ of each Speed Responsive Cell. Empty circles show the relationship of grating against narrow bandwidth MC, filled squares show grating against broad bandwidth MC. In both conditions most of the points fall below the line of identity. **f** Histograms show the distribution of *σ*_*v*_. Vertical red lines show the median of each distribution. The distribution for gratings is centered, while the distributions for both types of MC are skewed towards lower values with a significant difference (*T* = 1344 *p* = 0.00126 for grating vs. narrow bw. MC, *T* = 1013 *p* = 0.00001 for the broad bw. MC, Wilcoxon signed-rank test). **g,h** same as e,f but for the preferred speed (peak of the fitted curve) with a less significant difference (*T* = 1600 *p* = 0.026 for grating vs. narrow bw. MC, *T* = 2096 *p* = 0.731 for the broad bw. MC, Wilcoxon signed-rank test)

This change in the response results in tuning curves with narrower profiles, thus the tuning of the cell becomes more specific for certain parameters of the stimulus. Besides this change in the shape of the tuning curve, for many cells, the response to the MC stimuli has lower firing rates, as is the case for the example shown in Figure 5b, while for others, the responses are equivalent and even higher in some cases (see examples in Figure 5d).

Another interesting and unexpected finding is that some of the cells with narrower tuning for speed also present changes in their preferred speed, mostly towards higher speeds (Figure 5d, panels on the right).

To quantify the change in the tuning bandwidth of the grating versus MC response, we measured the properties of the curve fitted to the response to speed (see Methods section for details). In Figure 5d we show some examples of RGCs with narrower tuning in terms of lower response to the non-preferred speeds. At the population level, a large proportion of the SR cells show a decrease in the bandwidth of the response to speed at their preferred spatial frequency, for the complex stimuli when compared to the response to drifting gratings. This is evident when looking at the comparison of σ_*v*_ in Figure 5e; for both MC series, a large portion of the points falls below the line of identity, meaning that the values are lower than for the grating stimulus. The distributions of σ_*v*_ are skewed towards lower values (Figure 5f), meaning that a large portion of the speed responsive cells have narrower tunings when stimulated with the complex stimulus (*T* = 1344 *p* = 0.00126 for grating vs. narrow bw. MC, *T* = 1013 *p* = 0.00001 for the broad bw. MC, Wilcoxon signed-rank test). With regard to the preferred speed, even though that for some cells the preferred speed increases (see the clusters at the lowest and highest speeds in Figure 5g), overall, the differences are not highly significant (Figure 5h *T* = 1600 *p* = 0.024871 for grating vs. narrow bw. MC, *T* = 2096 *p* = 0.73165 for the broad bw. MC, Wilcoxon signed-rank test).

### Population responses becomes sparser for more naturalistic stimuli

When pooling the response of many cells, as a motion sensing neuron in higher cortical areas would do^19,40^, the response profile of the recorded population also shows differences when stimulated with the complex, more naturalistic stimuli. As expected from the individual responses, the response profile to speed depends on the spatial frequency (Figure 6a left panel). The curve not only shifts when changing the spatial frequency, but the peak response also decays as the spatial frequency increases.

**Figure 6.**
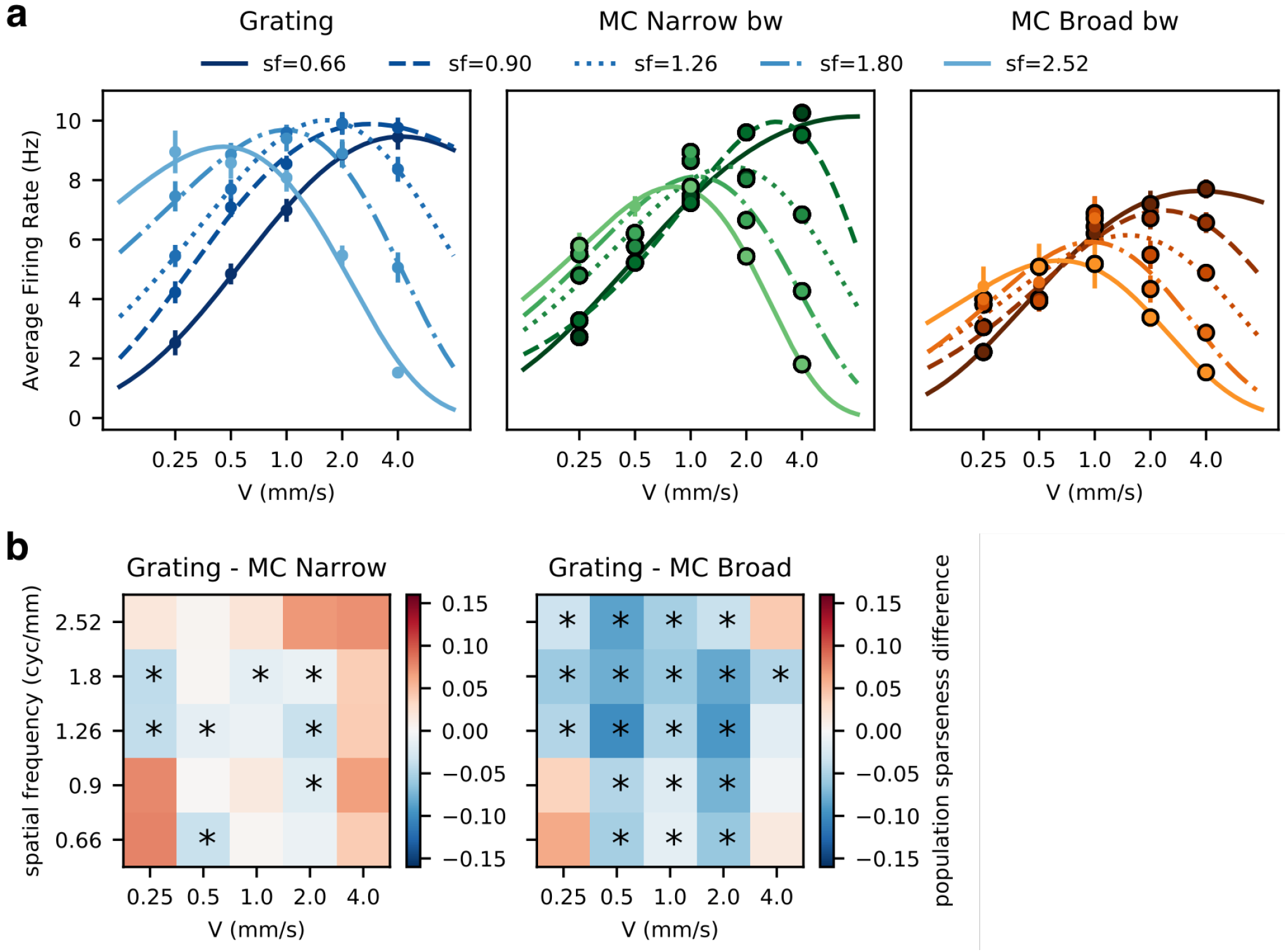
Naturalistic stimuli and sparse code. **a** Changes in Tuning to Speed at the population level. The dots show the average response across trials for the whole population of *Speed Responsive Cells* for each combination of speed and spatial frequency used; the lighter shades show increasing spatial frequency, the error bars show standard deviation. The encircled dots indicate all the conditions for which the response is significantly smaller than the response to the grating stimulus (*p* < 0.05 Wilcoxon signed-rank test). The response to speeds for each spatial frequency was fitted to an skewed Gaussian. As expected, at higher spatial frequencies the preferred speed shifts towards lower values, together with a decrease in maximum response, however, this decrease becomes more pronounced when the complexity of the stimulus increases, to the point where the response to lower speeds becomes very similar for different spatial frequencies (no significant difference at speeds 0.25 and 0.5 mms^−1^ (*p* > 0.05 Wilcoxon signed-rank test). **b** Sparseness in the code, calculated as population sparseness (Equation 4), is a measure of how many cells are responding and their relative level of activity. The 2-D plots show the average sparseness for each condition tested, for the three types of stimuli. In general, it is higher for the extreme parameters, however, for the more complex stimulus, the area of high sparseness is larger. The asterisks indicate the conditions for which the sparseness is higher compared to that obtained using grating stimulus (Wilcoxon signed-rank test, *p* < 0.05)

When comparing the response to the simple vs complex stimuli, the preference for lower speeds at higher spatial frequencies is still observed, but with a significant decrease in response for almost every condition (Figure 6a middle and right panels). However, this decrease in response is larger for the lower speeds at every spatial frequency tested, to the point that for the two lowest speeds, the difference in response between spatial frequencies becomes non-significant (no significant difference at speeds 0.25 and 0.5 mms^−1^ *p* > 0.05 Wilcoxon signed-rank test), so the left part of all the tuning curves converge to the same values.

However, these lower firing rates elicited by the complex stimuli would not necessarily be detrimental to the coding properties of the retina. Instead, lower average firing rates could be associated with sparse coding. To see if this were the case, we computed the average population sparseness (equation 4) for each stimulus condition (how many cells and how much they are responding to each stimulus). When compared to the simple stimulus, there are significant increases in sparseness in 8 condition for the MC with narrow bandwidth and 19 conditions for the broad bandwidth stimulus at each combination of speed and spatial frequency tested (Figure 6b).

### A sparse response encodes motion information efficiently

To determine if the sparser retinal response is still encoding the motion information, we applied a reconstruction framework to decode the trajectory of a moving stimulus from the response of a group of cells and then extract information derived from the trajectory. As can be seen in Figure 7, we constructed retinal images from the spikes emitted by each cell and its respective RF. To assess the ability to extract motion information from the reconstruction, we applied a process analogous to the way velocity is computed in cortical visual areas^41^ (see Methods section for details). An example of this process is shown in Figure 7. Even though that for some of the conditions the first stage of decoding was enough to get the correct speed (in the example, the lowest and the two highest speeds), in general the decoding improves at the Motion Energy and Opponent Motion stages, in the sense that the set of filters produces the highest activation is the one that matches the presented speed.

**Figure 7.**
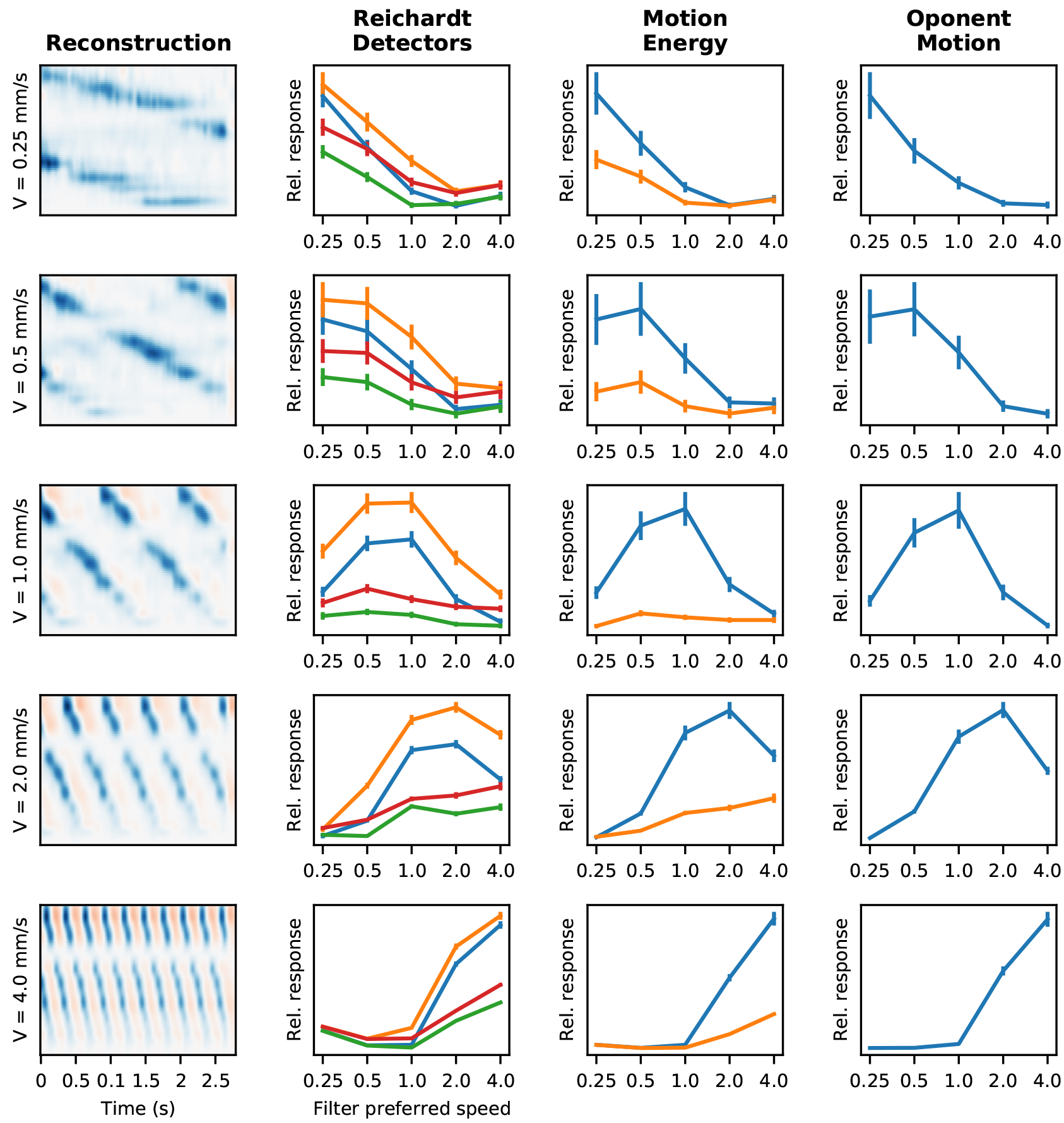
Reconstruction and decoding of motion stimuli. The leftmost column of each plot shows the reconstruction of the moving stimulus at different speeds, at a spatial frequency of 0.9 cycles/mm. From the population response, the stimulus is reconstructed by convolution of the spike train with the RF of each cell. The reconstructions are dominated by negative contrasts, due to the low number of ON type cells and their relatively low response. The mostly white row in each reconstruction corresponds to an area of the patch with no recorded units. The following columns show the response at each stage of decoding, for each corresponding reconstruction, averaged across trials. Vertical bars show s.d. The first stage is the response to each of the four filters that constitute the Reichardt detectors, the following column shows the Motion Energy in each direction and the rightmost column is the difference between the two. For each reconstruction the decoding is considered correct if, in the final stage, the preferred speed of the most activated filter matches the stimulus speed.

The precision of speed decoding for the different conditions can be seen in Figure 8. The motion traces obtained for all the stimuli can be seen in Figure 8a. To measure the quality of the estimation, we computed the error rate as 1 minus the number of trials for which the estimation is correct, over the total number of trials. As seen in Figure 8b, some of the conditions have zero error rate, while for the lowest speed, the error rate is 1 for many of the spatial frequencies for the three types of stimulus. For the gratings, most values are either one or zero, while for the MCs some conditions show intermediate values, meaning that for some trials the decoding was correct and for others it failed. When looking at the cumulative error rate across spatial frequencies (Figure 8c) it becomes apparent that the decoding error for gratings is very high at the lowest and highest spatial frequencies, decreasing gradually towards the intermediate spatial frequencies. The same pattern can be seen for the MCs, however, the difference in decoding error for different spatial frequencies is lower, and stays within an acceptable range for all conditions. For broad bandwidth stimuli, the error rate is distributed more evenly across spatial frequencies, with bad decoding performance concentrated at the lowest speed.

**Figure 8.**
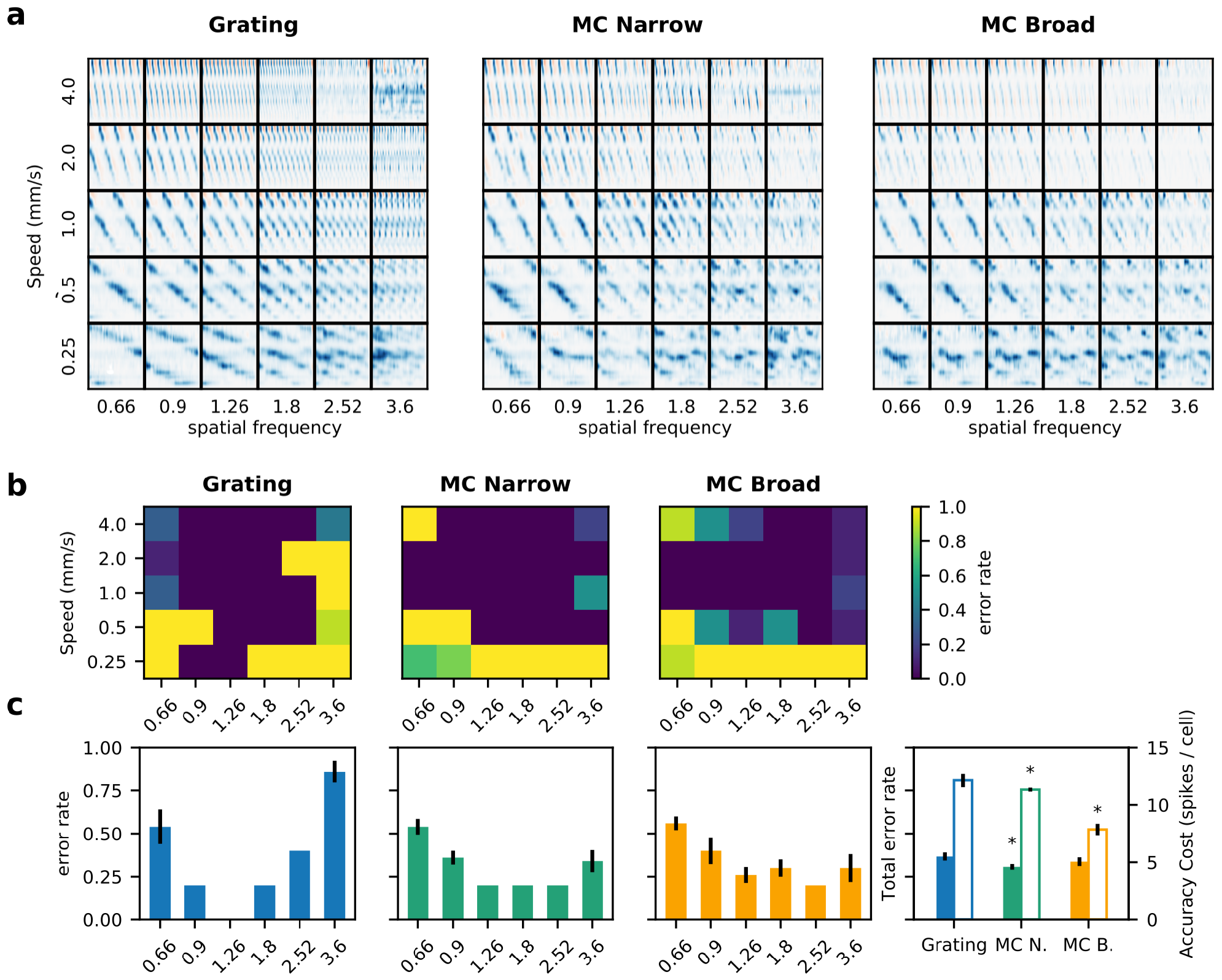
Precision of the estimation of stimulus speed. **a** Reconstruction of every motion stimulus tested. The representation is the same as in Figure 7. For the three types of stimulus, the motion traces (and their slopes) are easily seen for all conditions, except at the lower speeds and higher spatial frequencies, which is consistent with lower responses under those conditions (Figure 6). **b** Error rate in the decoding of the speed estimation (number of times that the algorithm estimated the correct speed over the total number of trials). The 2-D plots show the error rate at each spatial frequency and speed, for each type of stimulus. **c** Error rate for each spatial frequency and total error rate. Error lines show standard deviation between trials. In the rightmost plot, solid columns show total error rate and open bars show the Accuracy cost. Accuracy cost is computed as the success rate multiplied by the average firing (total number of spikes / number of cells); asterisks indicate significant difference with respect to the Grating (*p* < 0.05 Wilcoxon signed-rank test).

While the distribution of the errors is different for the three types of stimuli, overall, the total error rate is not very different between types of stimulus (Figure 8c, rightmost plot). The only significant difference is between gratings and MC of narrow bandwidth (*T* = 0.0, *p* = .0111 and *T* = 12.0, *p* = .205 Wilcoxon signed-rank test), but even there the magnitude of the difference is not large. Thus, it can be considered that the decoding performance is roughly equivalent. Since we were able to estimate the stimulus speed with an accuracy of ≈ 0.7 (1 - the total error rate) for the three types of stimuli, it could be argued from an information point of view it could be argued that the retina is transmitting at least enough information about the stimulus to encode speed under the three conditions. However, as shown in Figure 5, MCs elicit responses with lower firing rates. So, if we quantify the cost of accuracy for each type of stimulus, in terms of the number of spikes used to successfully transmit the speed of the stimulus, we get that for the gratings the value is ≈ 12.13 spikes/cell, while for the MC narrow and broad is 11.32 and 7.86 respectively, and these values are statistically significant (*T* = 7.0, *p* = .0366 for the MC narrow and *T* = 0.0, *p* = .0050 for the MC broad, Wilcoxon signed-rank test).

## Discussion

Our results strongly suggest that the tunning to natural stimuli is already present at the level of the retina. For the first time, we showed that the tuning curves of response to motion become narrower under stimulation with increasingly more complex images. In the present work, the difference between each set of stimuli (and thus the factor that would determine the differences in the response) is the spatial frequency content, *i.e*. the variety in sizes for the elements that constitute the images; while the three classes of stimuli have the same average properties (spatial frequency and speed) for each set, the MC contain additional signals or components around this central value.

Additional signals in an stimulus can modulate the response of a cell in different ways. It has been shown that stimuli that fail to modify the activity of a cell when presented in isolation, can modify the response to another stimulus when presented simultaneously^42^. This would stem from nonlinearities in the processing of the signal performed by the cell, in the sense that the response to a complex stimulus is not simply the response to the sum of its components. In some cases, the presence of signals around the RF, product of an increased area of stimulation, is enough to trigger changes in the response^21^. One of the mechanisms responsible for this effect is surround suppression, in which the presence of a stimulus in an area surrounding the spatial RF will decrease the response to the stimulus in the center^43^. This is partly explained by an increase in the amplitude of inhibitory post-synaptic potentials^22^.

In our experiments, all stimuli cover the same area, which exceeds the area of the RFs of the recorded cells, so the presence of stimulus in the surround would not by itself be enough to explain the changes in the observed response. Surround suppression occurs under a variety of stimulation patterns, both spatially and temporally^44^, however, for simple stimuli it has been shown that the effect is strongest when the parameters of the stimulus on the center and the surround are the same^45^, which would be the case for the gratings in our experiments; however, our data shows that this is not the case Figure 5.

Another factor to consider is the variance in contrast levels. Seminal works in vision psychophysics established that spatial frequency sensitivity is dependent on contrast and overall luminance levels^46,47^. In general, sensitivity to high spatial and temporal frequency decreases at lower contrast levels. It could be argued then that our results reflect the effects of changes in contrast and not changes in spatial frequency. In the proposed naturalistic stimulus, the relative contrast of each component of the images indeed scales with its spatial frequency, so components with higher spatial frequencies have lower contrast, following the same 1/*f* scaling characteristic of natural images. However, the contrast of the main components (those with the central spatial frequency sf_0_ or close to it) have the same contrast as the grating stimulus, while only the high frequency components have lower contrast, so the primary response drivers have the same contrast across the three types of stimuli. On the other hand, the effect of lower contrast on tuning curves is the opposite of what we report, as stimuli with lower contrast generate broader tuning curves^48,49^, at least in the visual cortex. Finally, it has been shown that adaptation can alter the relationship between spatiotemporal tuning curves and contrast^10^. Even though retinal adaptation to luminance levels is relatively fast (in the order of seconds), the stimulation time necessary to generate spatial frequency adaptation is longer, so we explicitly designed the stimulation protocol to avoid adaptation by limiting the exposure time to 3 seconds and interspersing the images with 1 second of blank, average luminance stimulation. The stability in the response during stimulation and across trials for the three classes of moving stimulus (Figure 5) shows that we can consider the effect of contrast, concerning the effect of spatial frequency bandwidth, to be negligible.

It has been shown that a non-linear interaction between the center and surround parts of the RF in RGCs modulates the response to spatial patterns in a context-dependent manner, and that this effect is greater during naturalistic stimulation^50^. We think that this effect can explain, at least in part, the results that we observe; the additional signals contained in the complex stimulus would act as non-linear inhibitors, resulting in lower response when the parameters are farther from the preference of the cell, producing narrower tuning curves.

Interestingly, together with this narrower bandwidth, approximately a third of the cells show an increase in preferred speed (see two examples in Figure 5, right panels), while others preserve their preference (Figure 5, left panels). The change in the speed preference cannot be attributed to changes in the spatial frequency content of the stimulus. Due to the bandwidth of the stimuli, the images will include additional spatial frequencies, but the shape of the spectrum (see Methods: Naturalistic stimuli) determines that most of the additional frequencies will be higher than the central frequency (sf_0_). If the cell processed these higher spatial frequencies linearly, they should shift preference towards lower speeds - but our results show the opposite. From our analysis, it is difficult to relate or predict this behavior from other properties of the cell’s response, since the shift in the preferred speed appears not tocorrelate with any of the RF’s characteristics.

Another point here is the implications or consequences of having these narrower tunings for naturalistic stimuli. At the level of the individual cell, it means that the cell will only respond when the stimulus is near the preferred parameters. But this does not mean that the cell will only encode its preferred stimulus. Indeed, it has been shown that steeper tuning curves will encode more efficiently because stimuli will be more easily discriminated in the high slope range ^51^. As such, this results means that for speed-selective cells, an input with a larger spectrum of spatial frequencies will have a higher precision in speed, *independently* of the local scale (spatial frequency) of the stimulus. This mechanism speaks to the dual mechanisms at the origin of selective responses (here speed) and of the invariance to other features (here scale). However, it is still not known how these features emerge in the low-level visual system, though computational studies predict that these properties emerge when using unsupervised learning to build models with a sparse coding constraint^13,52^. In addition to this effect at the individual cell level, when looking at the population code, narrower tuning curves can lead to less overlap between cells, so in conjunction the code will contain less redundancy, an important aspect of the efficient coding theory^53–56^. In support of this, we showed that population sparseness increases for the naturalistic stimuli, so in this respect the retina is performing better under these types of stimulation, in the sense that a sparse code is more efficient from an information transmission point of view ^57^.

To see if these changes in tuning curves have an effect on the coding of the motion signal, we implemented the stimulus reconstruction and speed decoding method. We chose speed as the relevant parameter for motion estimation, but we expect that with the proper adjustment, any property of the stimulus could be recovered, providing one is able to obtain a sufficiently good reconstruction^58^. Another simplification that we took advantage of, is that our experimental design is based on the independent presentation of a single speed for each stimulus set (with slight variations around an average value for the MCs); this means that for the duration of each segment of stimulation, the whole population of cells is “seeing” the same speed. Thus, to recover the speed of the stimulus from the reconstruction, we only need to get a single value from the whole population and for the whole presentation of each sequence.

Our reconstruction method could be improved by varying some of the parameters, such as the weights of the sum, to minimize the error of the results, as is usually done in this type of work. However, even though when the models obtained by those methods are biologically plausible, the process of *a posteriori* maximization of the quality of decoding is hardly naturalistic, because no biological sensory system has access to the “real” value of the signal to perform the minimization step that is essential to this kind of approach.

Another possible improvement would be to incorporate nonlinearities in the decoding process, possibly by taking into account the history of firing of a cell instead taking every spike with the same value. Nonlinear models have been reported to give better stimulus reconstruction, specially for complex images^59^.

In any case, even though our reconstructions have a relative low degree of fidelity, they contain enough information to successfully compute the speed of moving features in the images. The latter is important because it means that even though these complex stimuli elicit lower retinal responses, the relevant information is still contained in these lower responses. In this regime of lower firing rates, energy expenditure would be lower, and the cost of information would be lower^60^. Thus, this simplified approach is good enough to support the hypothesis that naturalistic stimuli would be encoded by the retina in an efficient way.

In conclusion, we have shown that the retina would be adapted to at least some of the characteristics of natural images, specifically those related to motion and the spatiotemporal information content and correlations. This adaptability is expressed in the narrower shapes of the tuning curves, which would lead to less overlap between cells in the feature space and thus less redundant and sparser population code, with smaller firing rates, which translates into less energy expenditure and less channel saturation.

## Methods

### Animals

*Octodon degus* born in captivity are maintained in a controlled facility. Prior to each experiment, animals were put in darkness for 30min, then deeply anesthetized with halothane and beheaded. Eyes were removed and dissected at room temperature under red illumination. Experimental procedures are approved by the Bioethics Committee and regulations from the University and in accordance with the bioethics regulation of the Chilean Research Council (CONICYT).

Octodon degus born in captivity are maintained in a controlled facility. Prior to each experiment, animals were put in darkness for 30 min, then deeply anesthetized with halothane and beheaded. Eyes were removed and dissected at room temperature under red illumination. Experimental procedures are approved by the Bioethics Committee and regulations from the University, and are in accordance with the bioethics regulation of the Chilean Research Council (CONICYT). Degus are diurnal rodents with 30% of their photoreceptors being cones and a comparatively high number of RGCs, qualities that makes them a good model for studying vision. In addition, the large area of the retina makes Degus especially suitable for Multi Electrode array recordings as good coverage of all the electrodes in the matrix can be obtained. For a complete description of the model see ^61,62^.

### Electrophysiological recordings

The experimental protocol was described before and follows^63^. Briefly, the physiological response of Degu RGC to different types of visual stimuli was measured using a Multi Electrode Array5^64,65^ (USB MEA256, Multichannel Systems GmbH, Reutlingen, Germany) using 256MEA100/30iR-ITO matrices and sampling at 20000Hz. After removing the eyes, the posterior hemisphere was dissected in quadrants and the pigmented epithelium separated from the retina. Finally, a piece of retina was mounted on a dialysis membrane ‘2and mounted on a perfusion chamber, which was then lowered onto the electrode array with the RGCs side down. Recording commenced under perfusion with AMES medium bubbled with 95% O_2_ 5% CO_2_ at 33 °C and the pH adjusted to 7.4.

### Visual Stimulus Presentation

Stimulus display was performed with a conventional DLP projector using custom optics to reduce and focus the image onto the photoreceptor layer, with size of a pixel of ≈4 μm, maintaining an average irradiance of 70nWmm^−2^. Stimuli were generated using the code available at https://github.com/NeuralEnsemble/MotionClouds. Timing of the images was controlled using a custom-built software based on Psychtoolbox for MATLAB^66^. Spike sorting analysis was performed using *Spyking-Circus*^67^. Data was analyzed via redistributable Jupyter notebooks using SciPy^68^ and running Python 3 kernels. All data fitting procedures were performed with LMFIT^69^. Figures were generated with Matplotlib^70^.

### Characterization of spatiotemporal tuning of RGCs

To produce a standard characterization of RGC responses to different stimulations, we measured the response to a white noise checkerboard pattern (block size = 0.05 mm) for 1200 s at 60 Hz and to sinusoidal drifting gratings at maximum contrast (minimum and maximum Weber contrast of ≈ −94 and ≈ 129 respectively), with spatial frequencies of 0.66, 0.9, 1.26, 1.8, 2.52 and 3.6 cycles/mm, and speeds of 0.25, 0.5, 1.0, 2.0 and 4.0 mm/ sec in sequences of three seconds with 10 repetitions of each. Due to the constraints in recording length stemming from tissue viability, we focused only on changes in response to the speed of the moving stimulus and not to its direction, so the present protocol uses a single direction for all sets of stimuli.

RC were estimated from the response to the checkerboard stimulus by reverse correlation, yielding the *Spike-Triggered Average (STA)*^71^ as a three-dimensional spatiotemporal impulse response (Figure 3E). The spatial characterization was performed by fitting a bi-dimensional Gaussian function at the time point of maximum amplitude, then drawing an ellipse at one standard deviation. The size of each RF was calculated as the radius of the circle with the same area as the ellipse fit^72^. The shape was quantified by the eccentricity ε of the ellipse as 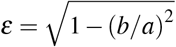 where *a* and *b* are respectively the radius of the major and minor axis; cells with an eccentricity larger than 0.9 were discarded. The temporal profile was computed as the time course of the intensity at the point with the largest variance, and then fitted to a difference of two cascades of low-pass filters^73^. Basal activity was set to zero and amplitude normalized, such that it follows the form

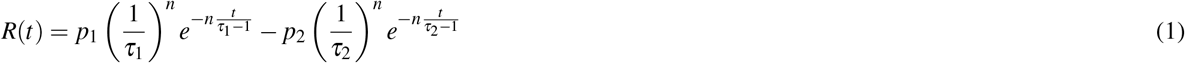

where *t* represents time in number of image frames before the spike, *τ*_1_ and *τ*_2_ describe the temporal decay of the response of each filter; scalars *p*_1_ and *p*_2_ are amplitude responses of each filter, and *n* is a free parameter. Tuning to motion was evaluated from the response to the drifting gratings; the response to each set of speeds and spatial frequencies was measured by *Peristimulus Time Histogram (PSTH)* for 10 repetitions of each sequence. The beginning of the PSTH is set to 200msec after stimulus start to discard the response to stimulus onset. Tuning to speed *v* was fitted to a skewed Gaussian in which the response to speed *R*(*v*) takes the form

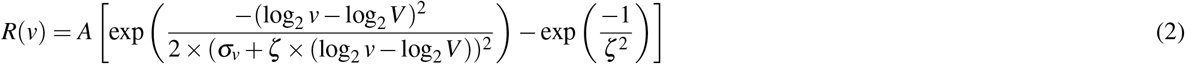

where *V* is the preferred speed, *σ*_*v*_ the curve width, ζ represents the skew of the curve and *A* a scaling factor^19^. Since *σ*_*v*_ is highly correlated with tuning bandwidth (width of the function at half-height)^74^, we used *σ*_*v*_ for all bandwidth related functions. Changes in tuning bandwidth were measured as the normalized difference in *σ*_*v*_ as

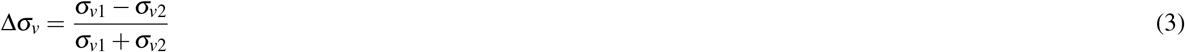

### Population coding

Population response was evaluated as the average of the PSTH of all *speed responsive cells*. Sparseness of the response to each stimulus was measured as Population Sparseness (*S*_*p*_), defined in^75^ as

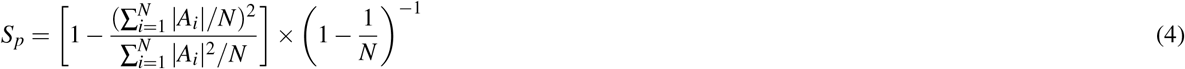

where *A*_*i*_ is the average response of cell *i* to the stimulus.

### Naturalistic stimuli

We used Motion Clouds (MCs)^37^ to measure the response of the retina to the motion of stimuli with different levels of naturalness (in terms of their spatio-temporal correlations). MCs are synthetic dynamical textures that mimic some key properties of natural images^38^, while allowing for the precise control over the signals that are being presented, which is a highly desirable property for the study of sensory systems under naturalistic conditions^17^. The traits of these textures are determined by three parameters for the center of the envelope (size, speed, and orientation), along with their respective bandwidths. These bandwidths control the area (in parameter space) in which the power of the stimulus will be spread.

Our protocols consisted of a set of different sequences with the same motion information (mean speed and spatial frequency), but in which we progressively varied the spatial frequency bandwidth (*i.e*. the range of sizes), as this was reported to influence visual motion detection^23^. These match the parameters of the drifting grating stimuli (equivalent to an MC with infinitely narrow bandwidth), and textures with a narrow and a wide bandwidth of spatial frequencies (Figure 1a blue, green and orange ellipses respectively). For narrow and broad bandwidth textures, amplitude scales with spatial frequency: for the narrow bandwidth stimuli, the scaling factor is chosen for the spectra to not overlap, while for the broad band stimuli the scaling factor is 1 resulting in *B*_sf_ = sf_0_, so that each spectrum overlaps with the adjacent one along the line of equal speed, see table 1 for the specific parameters. Figure 1b shows an example stimulus, comparing the spatiotemporal image of a drifting grating and MCs with narrow and broad bandwidth, illustrating the smooth transition from a “cristal-like” pattern (the grating) to progressively more complex and naturalistic stimuli using a single parameter (*B*_sf_). Under this characterization, a collection of well-sampled natural images would correspond to MCs with infinite bandwidth, *i.e*. it would contain every spatial frequency, so drifting gratings and natural images would be on opposite ends of the complexity scale, with our MCs constituting a gradual transition between them.

**Table 1.**
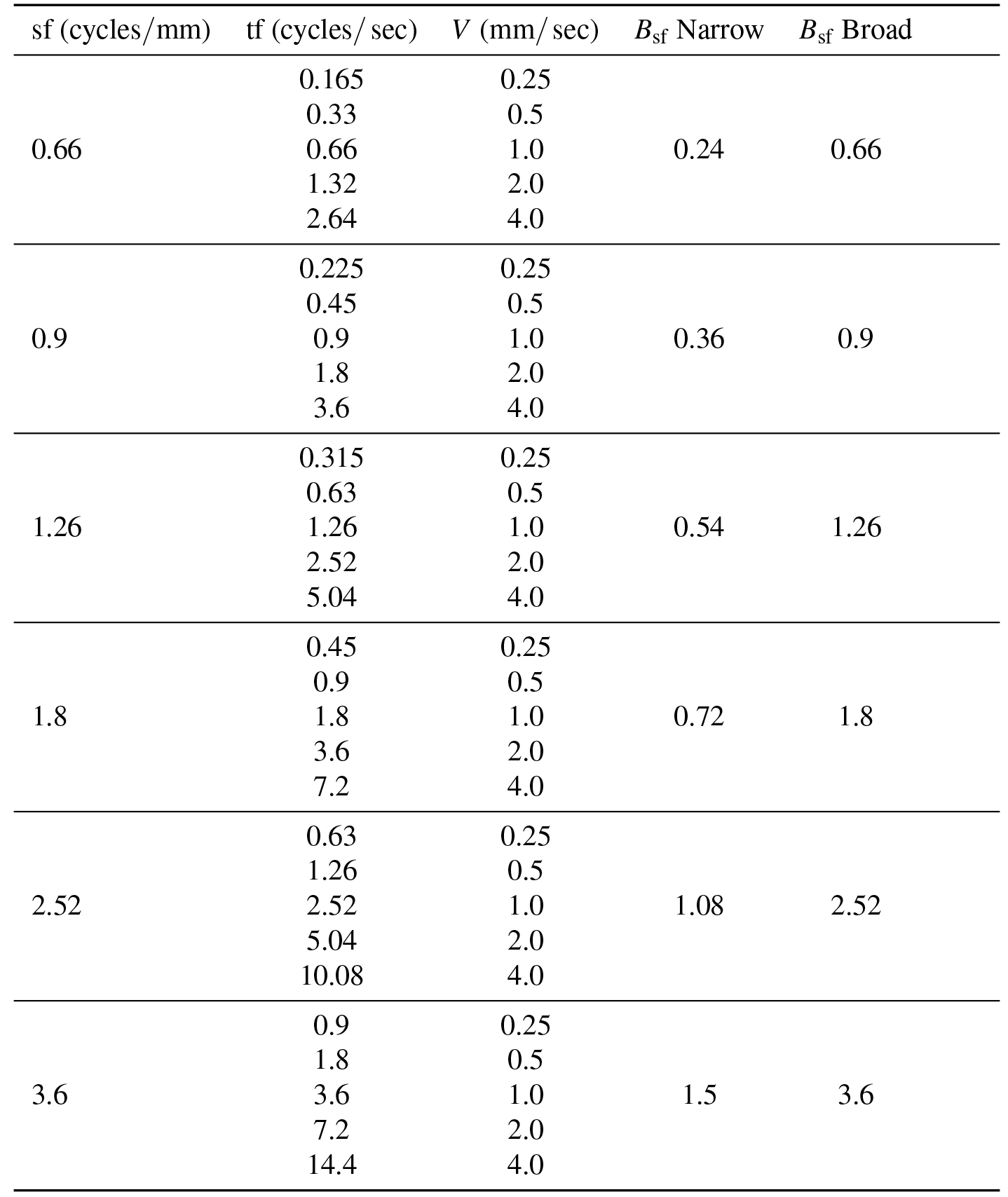
Parameters used to build each set of motion stimuli. For the gratings, the temporal frequency tf is modulated to obtain the same speeds at every spatial frequency sf. Motion Clouds do not have a tf parameter, they are controlled directly by the speed parameter *V*. Note that the Motion Clouds software works in pixel units, so a proper scaling has to be applied. In our case, the conversion factor between pixels and mm is 240:1

| sf (cycles/mm) | tf (cycles/sec) | V (mm/sec) | $B_{sf}$ Narrow | $B_{sf}$ Broad |
| --- | --- | --- | --- | --- |
| 0.66 | 0.165 | 0.25 | 0.24 | 0.66 |
|  | 0.33 | 0.5 |  |  |
|  | 0.66 | 1.0 |  |  |
|  | 1.32 | 2.0 |  |  |
|  | 2.64 | 4.0 |  |  |
| 0.9 | 0.225 | 0.25 | 0.36 | 0.9 |
|  | 0.45 | 0.5 |  |  |
|  | 0.9 | 1.0 |  |  |
|  | 1.8 | 2.0 |  |  |
|  | 3.6 | 4.0 |  |  |
| 1.26 | 0.315 | 0.25 | 0.54 | 1.26 |
|  | 0.63 | 0.5 |  |  |
|  | 1.26 | 1.0 |  |  |
|  | 2.52 | 2.0 |  |  |
|  | 5.04 | 4.0 |  |  |
| 1.8 | 0.45 | 0.25 | 0.72 | 1.8 |
|  | 0.9 | 0.5 |  |  |
|  | 1.8 | 1.0 |  |  |
|  | 3.6 | 2.0 |  |  |
|  | 7.2 | 4.0 |  |  |
| 2.52 | 0.63 | 0.25 | 1.08 | 2.52 |
|  | 1.26 | 0.5 |  |  |
|  | 2.52 | 1.0 |  |  |
|  | 5.04 | 2.0 |  |  |
|  | 10.08 | 4.0 |  |  |
| 3.6 | 0.9 | 0.25 | 1.5 | 3.6 |
|  | 1.8 | 0.5 |  |  |
|  | 3.6 | 1.0 |  |  |
|  | 7.2 | 2.0 |  |  |
|  | 14.4 | 4.0 |  |  |

### Trajectory Reconstruction and velocity estimation

The first step is the reconstruction of the trajectory of a moving stimulus from the response of a population of RGCs. As recently demonstrated in^59^, the *Decoding Fields* obtained by maximum likelihood are very similar to the RFs obtained by traditional methods of reverse correlation. Based on this, we made a spatiotemporal reconstruction of the stimulus by convolving the response vectors of each cell with its respective RF and then performing a weighted sum across the whole population. The response 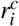 is the number of spikes at each interval *i* with Δ*t* = 16.66 ms. The RF is computed by reverse correlation from the response to a checkerboard stimulus. Since we are interested in stimuli moving only in one direction, we collapse the tridimensional spatiotemporal RF to a bidimensional representation 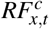 containing only time and one spatial dimension. Thus, the intensity of the reconstructed stimulus at each point in space and time, *I*(*x*, *i*), is given by

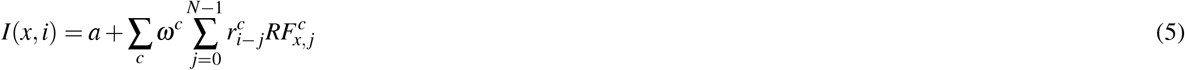

where *a* is a constant offset, *ω*^*c*^ is the weighting factor for each cell, 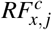 is the value of the receptive field at point *x* at time *j*Δ*t* before the spike, and *N*Δ*t* is the length of the RF estimation.

Speed of the moving stimulus is assessed by a multi-stage process following standard motion processing models^41^. The first step is the convolution of the reconstructed stimulus with sets of filters resembling Reichardt detectors, approximated by slanted Gabor filters oriented in space time with preference for different speeds (given by the slope in the space/time plane, thus the speed tuning is given by the angle of the Gabor kernel). Each kernel is paired with another of the same properties, but with its phase shifted, forming a quadrature pair. The response of each quadrature pair is rectified and added to obtain the Motion Energy for each speed. The process is repeated to compute the Motion Energy in the opposite direction, and finally, both are subtracted to obtain the net motion signal for each speed. The estimated speed is determined by the set of filters with the highest activation in a winner-takes-all approach. An example of the process is shown in Figure 7.

## Acknowledgements

Fondecyt 1150638 and 1140403, CONICYT-Basal Project FB0008, Millennium Institute ICM-P09-022-F, L.U.P. was supported by ANR projects “BalaV1” (ANR-13-BSV4-0014-02) and “TRAJECTORY” (ANR-15-CE37-0011). C.R. was supported by a CONICYT Ph.D. scholarship.

## Author contributions statement

M.J.E and A.P. conceived the study. A.P. provided the animals, equipment for electrophysiology and data analysis. C.R. designed the stimulation protocols and conducted the experiments. C.R. and L.P. analyzed the results. C.R. and M.J.E. prepared the figures. All authors reviewed the manuscript.

## Additional information

**Competing financial interests** the authors declare no competing interests.

